# Sperm derived H2AK119ub1 is required for embryonic development in *Xenopus laevis*

**DOI:** 10.1101/2024.04.23.590676

**Authors:** Valentin Francois--Campion, Florian Berger, Mami Oikawa, Maissa Goumeidane, Nolwenn Mouniée, Vanessa Chenouard, Kseniya Petrova, Jose G Abreu, Cynthia Fourgeux, Jeremie Poschmann, Leonid Peshkin, Romain Gibeaux, Jérôme Jullien

## Abstract

Deposition of H2AK119ub1 by the polycomb repressive complexe-1 plays a key role in the initiation of facultative heterochromatin formation in somatic cells. Here we evaluate the contribution of sperm derived H2AK119ub1 to embryo development. In *Xenopus laevis* we found that H2AK119ub1 is present during spermiogenesis and into early embryonic development, highlighting its credential for a role in the transmission of epigenetic information from the sperm to the embryo. *In vitro* treatment of sperm with USP21, a H2AK119ub1 deubiquitylase, just prior to injection to egg, results in developmental defects associated with gene upregulation. Sperm H2AK119ub1 editing disrupts egg factor mediated paternal chromatin remodelling processes. It leads to post-replication accumulation of H2AK119ub1 on repeat element of the genome instead of CpG islands. This shift in post-replication H2AK119ub1 distribution triggered by sperm epigenome editing entails a loss of H2AK119ub1 from genes misregulated in embryos derived from USP21 treated sperm. We conclude that sperm derived H2AK119ub1 instructs egg factor mediated epigenetic remodelling of paternal chromatin and is required for embryonic development.

## INTRODUCTION

Several studies point towards an epigenetic contribution of sperm for embryonic development. First, in several species paternal exposure to various environmental insults results in changes to the offspring ^1–5^. In addition to such environmentally induced epigenetic changes, sperm is also proposed to naturally harbour epigenetic cues that are required for development. Histones have been considered as possible vectors for such epigenetic information. Indeed, in several species including human, modified histones such as H3K4me3 and H3K27me3 are found in the sperm chromatin, in particular around developmental genes and in a pattern reminiscent of that found in embryonic stem cells^6–9^. In a previous work we provided evidence that in the frog, *Xenopus laevis,* some of these modified histones are homogenously distributed in a population of sperm cells, hence indicating that they have the required attributes for a role in the regulation of embryonic development^10^. Lastly, interference with histone variant or post-translational modifications during the formation of the gametes, or after fertilisation, lead to embryonic gene misregulation^9,11,12^. Although these later experiments support a functional role for sperm-derived modified histone in embryonic development, indirect effect of the interference on the process of spermiogenesis or early embryos development cannot be ruled out. A direct assay to test for the requirement of sperm modified histones in embryo development is still lacking. This is due in part to the difficulty to access the chromatin in the highly compacted sperm nucleus. In this work, we aim to fill this gap and devised a strategy to alter the sperm epigenome just prior to its introduction to the egg. We specifically considered H2AK119ub1 as a possible carrier of necessary epigenetic information from the sperm to the embryos. Indeed, recent works points towards a key role of polycomb repressive complexe-1 (PRC1), upstream of PRC2, in the establishment of facultative heterochromatin, both in cultured cells and in embryos^13^. The H2AK119 ubiquitylation activity of the PRC1 complex appears to be integral part of this mechanism^13,14^. In *Xenopus laevis* we found that H2AK119ub1 dynamic during spermiogenesis and early embryonic development is compatible with a role in the epigenetic programming of sperm for embryonic development. We applied an *in vitro* epigenetic editing assay that uses USP21 deubiquitylase activity ^15–18^ to erase H2AK119ub1 from sperm chromatin. Embryos derived from such epigenetically edited sperm are developmentally impaired. Transcriptome and epigenome analysis indicates that H2AK119ub1, in combination with H3K4me3, is enriched on a set of embryonic genes misregulated in embryos derived from USP21 treated sperm. Such USP21 sensitive genes preferentially retain H2AK119ub1 upon sperm chromatin exposure to maternal factors. Lastly, we provide evidence that egg factor mediated deposition of H2AK119ub1 on paternal chromatin is influenced by pre-existing sperm H2AK119ub1.We conclude that sperm derived H2AK119ub1 instructs egg factor mediated epigenetic remodelling of paternal chromatin and is required for embryonic development.

## RESULTS

We first evaluated the overall levels of H2AK119ub1 in chromatin during spermiogenesis and early embryonic development. Indeed, the presence of such modified histone in sperm seems to differ between species, its presence documented in that of mouse^19^ while it appears absent in that of zebrafish^20^. In frogs the last stage of spermatogenesis is associated with a remodelling of nucleosomes resulting in a partial loss of H2A/H2B while a full complement of H3/H4 is retained on sperm chromatin (**Fig. 1A**)^10^. Western Blot (WB) analysis captures this loss of H2A and reveal an overall decrease in H2AK119ub1 in sperm compared to spermatid (**Fig. 1B and Suppl Fig.S1-A**, see H2A/H4 and H2AK119ub1/H4 ratio, respectively). Importantly however, the H2AK119ub1 to H2A ratio is similar in sperm compared to spermatid, indicating that in this context of spermiogenesis-associated H2A loss, a significant level of H2AK119ub1 is maintained in sperm (H2AK119ub1/H2A, **Fig. 1B**). We next assessed H2AK119ub1 occurrence after fertilisation: in early frog embryos around ZGA (blastulae stage) and later during gastrulation and neurulation. Interestingly, contrary to what is seen with H3K27me3, a PRC2 associated modification, H2AK119ub1 is detected throughout early embryogenesis (**Fig. 1C**). Altogether, these observations highlight H2AK119ub1 credentials for a potential role in the transmission of epigenetic information from sperm to the embryos.

**Figure 1:**
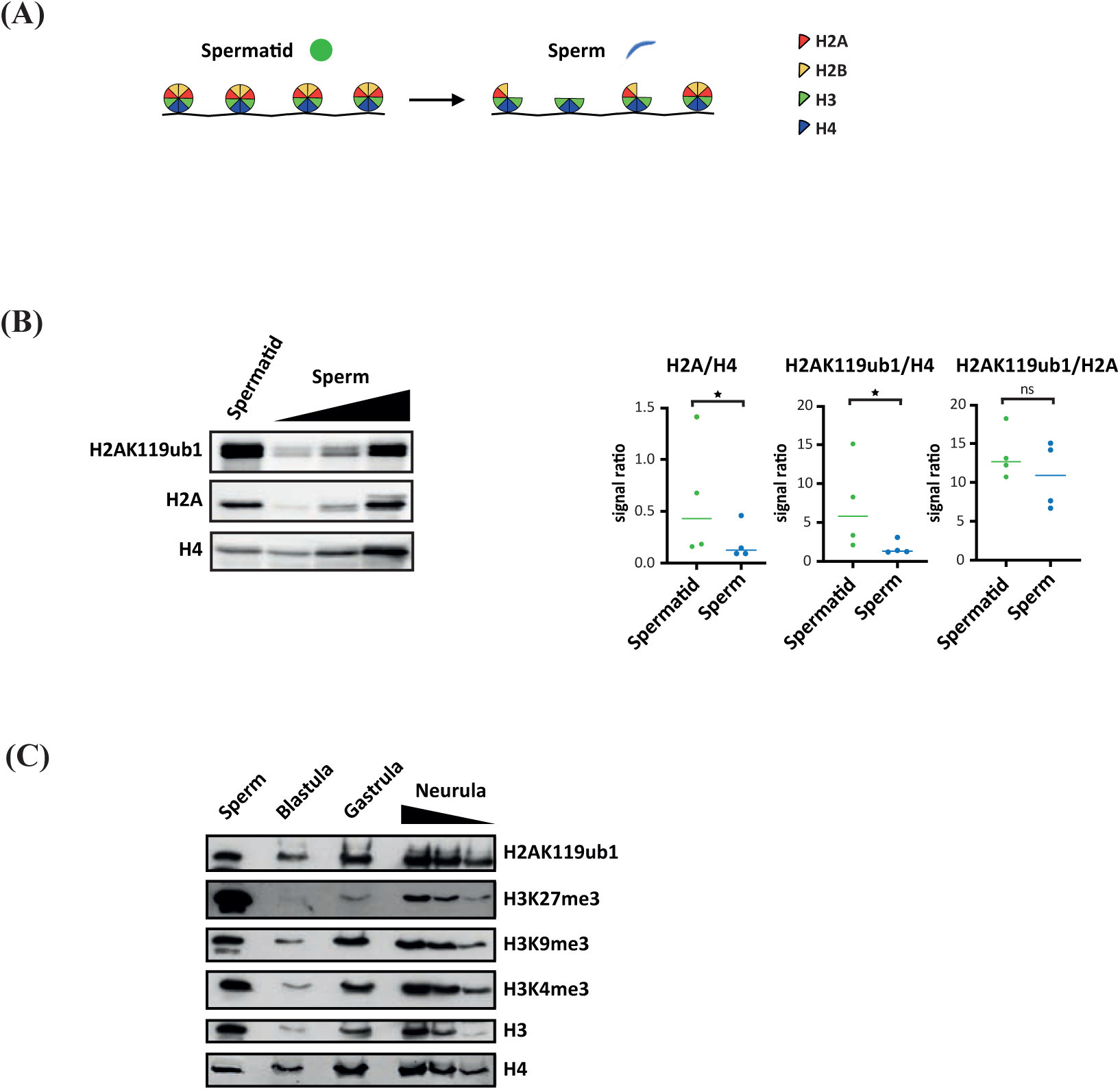
H2AK119ub1 dynamics during spermiogenesis and early embryonic development. (**A**) Schematic representation of *Xenopus laevis* nucleosome remodelling during spermiogenesis (**B**) Left, Western blot analysis of H2AK119ub1, H2A and H4 histones. Right quantitation of signal ratio of H2A to H4, H2AK119ub1 to H4, and H2AK119ub1 to H2A, in spermatid and sperm. * p<0.05, ratio paired t test (n=4). Median is shown. (**C**) Western blot analysis of H2AK119ub1, H3K27me3, H3K9me3, H3K4me3, H3 and H4 in *Xenopus laevis* embryo pre- (blastulae) and post- (gastrulae, and neurula) ZGA.

We then carried out H2AK119ub1 ChIP to map the distribution of this histone post-translational modification (PTM) on the sperm genome. Biological triplicates were well correlated and subsequently pooled for downstream analysis (**Suppl Fig. 2A-D**). We first assessed the distribution of H2AK119ub1 around the transcription start of all sperm genes. H2AK119ub1 accumulates around the TSS of a fraction of sperm genes (**Fig. 2A**), similar to what is observed in somatic cells^21^ and embryos^14,20^. Noticeably, we observed in the top 10% TSS harbouring H2AK119ub1 an enrichment for genes associated with developmental function (**Fig. 2A, Suppl Table 1**), and reminiscent of the association reported with H3K27me3 in many species^6–9^. We next sought to more generally identify regions associated with H2AK119ub1 genome wide. H2AK119ub1 is a pervasive histone modification typically present on ∼10% of nuclear H2A in somatic cell^22,23^ and up to ∼40% in germ cells^19^. In *Xenopus laevis* sperm, it accumulates over region covering larger part of the genome than other modified histones with broad distribution such as H3K27me3 (**Suppl Fig. 2E**). To identify region of H2AK119ub1 genomic enrichment we therefore used RECOGNICER, a coarse-graining approach designed to capture enrichment over broad scale^24^ (**Suppl Fig. 3, Table S1**). Although we observed peaks of accumulation around genes, such as illustrated for *wnt1.L*, *pax6.L* or the *hoxa.S* cluster (**Fig. 2B)**, a large proportion of H2AK119ub1 also accumulates on the sperm genome in distal region away from genes (**Fig. 2C**). Noticeably we observed that H2AK119ub1 is significantly associated with sperm gene regulatory elements (promoters and enhancers, **Fig. 2D**). Epigenetic control of gene expression often entails a combination of epigenetic cues such as those observed in X inactivation^25^ or in transcriptional reprogramming^26^. We therefore investigated the relationship between genomic regions enriched in sperm for H2AK119ub1 and for other histone post-translational modifications. We focused on H3K27me3^10^, associated with the activity of Polycomb Repressive Complex 2, and H3K4me3^10^, a histone modification associated with transcription (Peaks, *e.g.* around *pax6.L* and *hoxa.S* loci, **Fig. 2B**). We then determined the genome-wide association of H2AK119ub1 with methylated K4 and/or K27 on histone H3. While most H2AK119ub1 peaks exist independently of these methylated histones, ∼24% of H2AK119ub1 peaks overlap with either H3K4me3 and/or H3K27me3 peaks (**Fig. 2E**). We observed a very strong promoter bias for peaks of H2AK119ub1 co-occurring with H3K4me3, an expected trend given that H3K4me3 is known to strongly associate with gene promoter (**Fig. 2F**). By contrast, when occurring in the absence of H3K4me3, H2AK119ub1 is located away from promoter and in intergenic regions. Analysis of fragment length obtained following sequencing of ChIP and Input fractions allow the discrimination of DNA fragment associated with nucleosomes (full complement of histones protecting 150bp of DNA) from those associated with subnucleosomes (nucleosomes lacking one or two H2A-H2B dimer and protecting 110 or 70bp of DNA) in sperm chromatin (**Suppl Fig. 4**). Noticeably, in sperm chromatin on the whole, H2AK119ub1 is mostly associated to nucleosomes while H3K27me3 and H3K4me3 tend to be associated with subnucleosomes. So, while we observe co-enrichment of H2AK119ub1 with methylated histone H3 at a subset of genomic loci, those histone modifications are possibly present at those sites on adjacent nucleosomal/subnucleosomal nucleoprotein complexes rather than on the same particle. Altogether, this survey of H2AK119ub1 distribution in sperm reveals its enrichment on gene regulatory region further strengthening its potential for gene regulation across generation.

**Figure 2:**
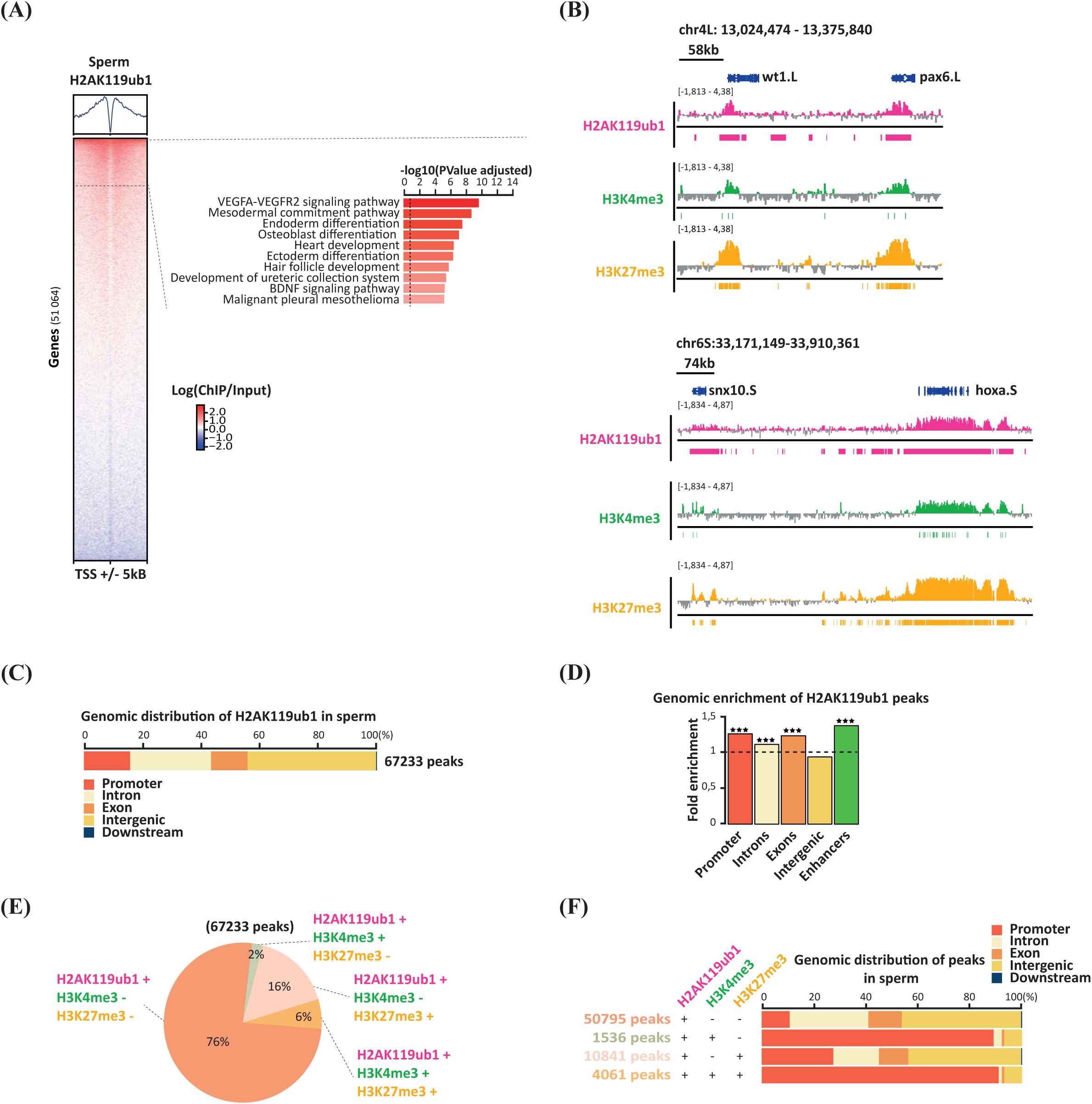
H2AK119ub1 associates with gene regulatory regions in sperm chromatin. **(A)** Heatmap of H2AK119ub1 signal (log(ChIP/Input)) on all *Xenopus laevis* genes TSS +/-5kB. GO enrichment is shown for the top 10% genes. **(B)** Genome browser view of log(ChIP/Input) of sperm H2AK119ub1, H3K4me3, and H3K27me3 around *wt1.L, pax6.L*, *snx10.s* and *hoxa.S* loci. Boxed area underneath the gene track indicates regions of enrichment (peaks). **(C)** Distribution of sperm H2AK119ub1 peaks on genomic features. **(D)** Fold enrichment of H2AK119Ub1 peaks on different genomic features, *** p<0.001, permutation test. **(E)** Pie chart indicating the proportion of H2AK119ub1 peaks either alone, or overlapping H3K4me3 and/or H3K27me3 peaks in sperm. **(F)** Genomic distribution of sperm H2AK119ub1 peaks according to overlap with H3K4me3 and/or H3K27me3 peaks.

To further evaluate how sperm H2AK119ub1 could have an impact on embryonic development, we next sought to follow its fate after fertilisation. Following delivery to the egg, protamines are removed from sperm chromatin and a canonical nucleosomal structure is re-instated. In conjunction with these events the sperm chromatin is replicated. Therefore, the extensive changes to sperm chromatin that follow fertilization have the potential to profoundly alter the sperm derived modified histone landscape. To determine the fate of sperm modified histones we turned to interphase egg extracts that can mimic the sperm chromatin remodelling events that follow fertilization, including protamine to histone transition and DNA replication^10^. We carried out H2AK119ub1 and H3K4me3 ChIP-seq on untreated sperm, or sperm having gone through egg extract mediated replication (**Fig. 3A**). Combining this dataset with existing spermatid and gastrulae ChIP-seq^9,10,27^ we obtain a global view of H2AK119ub1, H3K4me3, and H3K27me3 dynamic during spermiogenesis and early embryogenesis (**Fig. 3B**). Remarkably, while H3K4me3 and H3K27me3 are mostly stable, H2AK119ub1 exhibits drastic redistribution throughout this developmental period. Indeed, a large set of genomic location acquires H2AK119ub1 following sperm chromatin replication (cluster 4, **Fig. 3B, Suppl Table 2**), and includes a large proportion of gene TSSs (**Suppl Fig. 5A**). To better delineate the principle of this developmental dynamic, we then compared H2AK119ub1 and H3K4me3 peaks in untreated (sperm) *versus* egg extract treated sperm (replicated sperm). We confirm a marked disparity in replication dynamics between sperm H3K4me3, mostly stable (77% of sperm peaks conserved after replication (19345/24982) and 52% stable peaks overall (19345/36770)), and sperm H2AK119ub1, mostly changing (23.5% of sperm peaks maintained after replication (15796/67233) and 15% stable peaks overall (15796/108740)) (**Fig. 3C, Suppl Fig. 5B)**. Most of the replication redistribution of H2AK119ub1 happens in intergenic region, while H2AK119ub1 peaks stable through this process are mostly located on promoter (**Suppl Fig. 5D**). In this dynamic landscape, the subset of H2AK119ub1 genomic location associated with H3K4me3 in sperm appears selectively conserved (93% of sperm bivalent peaks are maintained after replication (5222/5597) (**Fig. 3C**). Broader H2AK119ub1 peaks with higher level of H2A retention discriminate location where H2AK119ub1/ H3K4me3 is maintained following exposure to egg factors from those where it is lost (**Suppl Fig. 5C**). Lastly, we confirmed that the H2AK119ub1 configuration observed following *in vitro* treatment of sperm with egg extract is also detected in blastulae stage embryo at the time of ZGA (**Suppl Fig. 6A-C**). We conclude that sperm exposure to egg factors profoundly remodels paternal H2AK119ub1 genomic distribution.

**Figure 3:**
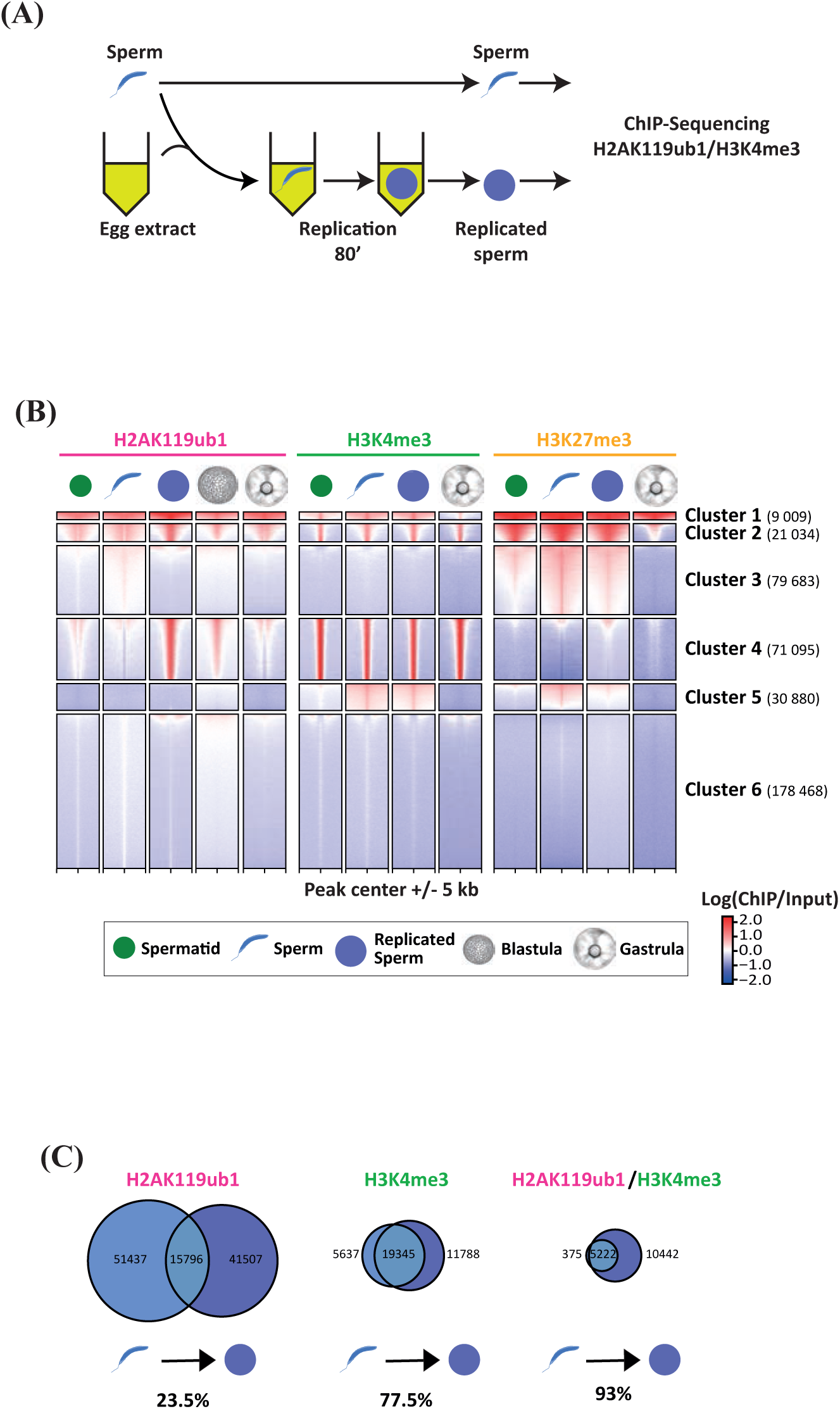
Sperm H2AK119ub1 is remodelled following exposure to egg factors. **(A)** Experimental design for monitoring modified histones fate following sperm exposure to egg factors. (**B**) Heatmap of H2AK119ub1, H3K4me3, or H3K27me3 signal (log(ChIP/Input)) in spermatid, sperm, replicated sperm, and gastrula. All genomic locations enriched for one of these histone modifications (peaks) in at least one of these developmental stages are represented. **(C)** Venn diagram indicating the changes to H2AK119ub1 (left), H3K4me3 (middle) or overlapping H3K4me3/H2AK119ub1(right) peaks during sperm chromatin replication in egg extract. Percentages of sperm peaks that are maintained after replication are indicated underneath.

We next evaluated to which extent such egg factor-mediated H2AK119ub1 remodelling is dependent on incoming sperm chromatin component. To directly evaluate sperm H2AK119ub1 functional relevance to this process, we set out to erase this histone modification, after completion of spermatogenesis, and prior to exposure to egg factors. To that end, we incubated mature sperm in an oocyte extract containing USP21, a H2A deubiquitylase^15,28^ (**Fig. 4A**). In such extract, the sperm nucleus is partially decondensed, allowing efficient deubiquitylation of H2AK119ub1 while the levels of trimethylated H3K4, H3K9 and H3K27 remain unchanged (**Fig. 4B, Suppl Fig. 7A, Suppl Fig. 1B**). Importantly, MNAse digestion of oocyte extract USP21-treated sperm yields nucleosomal and subnucleosomal size DNA fragments typical to that of naïve *Xenopus* sperm^9^ (**Fig. 4C**, **Suppl Fig. 7B, and 7C left**). This indicates that the overall sperm chromatin structure is maintained after oocyte extract treatment and does not convert to a somatic type of chromatin organisation as triggered by egg extract treatment^29^ (**Fig. 4C, Suppl Fig. 7**). We next asked to what extend USP21 mediated erasure of H2AK119ub1 from sperm chromatin influences its egg factor mediated redistribution. To that end, we carried out modified histone analysis in sperm treated with control or USP21 extract oocyte with or without a subsequent incubation in egg extract (**Suppl Fig. 8A**). Interestingly, WB analysis indicates that H2AK119ub1, erased from sperm chromatin by USP21 treatment, is at least partially reinstated by subsequent exposure to egg extract (**Fig. 4D**), indicating the existence of a maternal H2AK119 ubiquitylation activity independent of pre-existing sperm H2AK119ub1. Using ChIP-seq, we next ask if sperm H2AK119ub1 erasure prior to exposure to egg components affects the maternal factor mediated remodelling of H2AK119ub1 on paternal chromatin. We first focused on sperm genomic sites where we previously detected accumulation of H3K4me3, H3K27me3, or H2AK119ub1 (as in **Fig. 3B)**. This analysis confirmed that USP21 efficiently erases H2AK119ub1 from its genomic site in sperm (**Suppl Fig. 8B, Suppl Table 3**). We observed an overall reduction in egg factor mediated H2AK119ub1 deposition on chromatin derived from USP21+ sperm compared to that from USP21-sperm (**Suppl Fig. 8B**). We then compared H2AK119ub1 peaks in samples originating from replicated USP21-treated sperm to that from replicated control-treated sperm (**Fig. 4E**). Surprisingly, we observed that sperm USP21 treatment leads not only to loss of H2AK119ub1 peaks post-replication (lost peaks) but also triggers the appearance of a large set of peaks that is not deposited by maternal factors in control sperm samples (gained peaks) (**Fig. 4E**). Comparatively, USP21 treatment impact on post-replication H3K4me3 is much milder (**Suppl Fig. 8C**). To better characterize this shift, we first evaluated if H2AK119 ubiquitylation sites detected following egg extract exposure of sperm harbour genomic features associated with variant PRC1 (vPRC1) targeting in somatic cell (reviewed in^30^). Those H2AK119ub1 sites that are specific to egg extract treated control sperm (lost peaks, **Fig. 4E**) are associated with cluster 4 in Fig.3B and are indeed enriched for TF motifs associated with vPRC1 targeting (*i.e.* USF1/2 E2F, E-box, **Suppl Fig. 8D, Suppl Table 2**). We also detected a strong association of these sites with unmethylated CpG island (**Fig. 4F-G**), a genomic feature also shown to attract vPRC1 activity. By contrast, sperm treatment with USP21 prior to egg factor exposure redirects H2AK119ub1 accumulation towards repeat elements and away from such targets (**Fig. 4H-I, Suppl Fig. 9, Suppl Table 5**). We conclude that maternal factors mediated redistribution of sperm H2AK119ub1 is instructed by pre-existing H2AK119ub1 derived from paternal chromatin. In the absence of sperm H2AK119ub1, maternal factor mediated H2A ubiquitylation activity is shifted from CpG islands to repeat elements.

**Figure 4:**
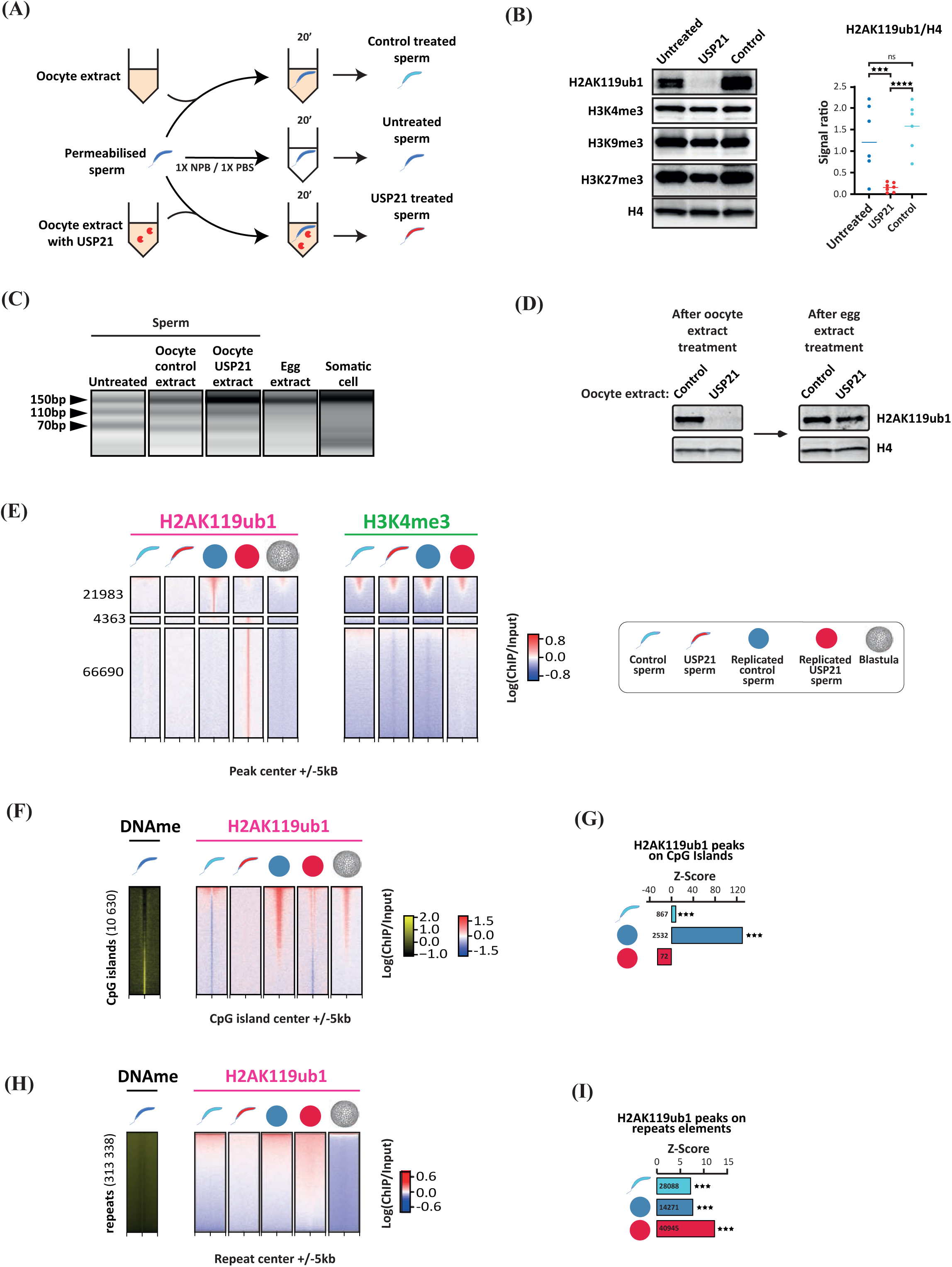
H2AK119ub1 remodelling by egg factors is altered by sperm pre-treatment with a H2A deubiquitylase. **(A)** Schematic representation of sperm treatment with oocyte extract containing USP21. **(B)** Left, Western blot of H2AK119ub1, H3K4me3, H3K9me3, H3K27me3 and histone H4 on untreated sperm, USP21 treated sperm and oocyte extract treated sperm. Right, signal ratio of H2AK119ub1 to H4 in untreated sperm, USP21 treated sperm, or oocyte extract treated sperm. *** p<0.005, **** p<0.001, ratio paired t test (n=6). Median is shown. (**C**) DNA fragments generated by MNase digestion of *Xenopus laevis* untreated sperm, oocyte control extract treated sperm, oocyte USP21 extract treated sperm, egg extract treated sperm and somatic cell. (**D**) Western blot analysis of H2AK119ub1 and histone H4 on sperm after treatment with control- or USP21- oocyte extract without (left) or with (right) subsequent replication in egg extract. (**E**) Heatmap of H2AK119ub1 and H3K4me3 signal (log(ChIP/Input)) in sperm after treatment with control- or USP21- oocyte extract without or with subsequent replication in egg extract. All genomic locations enriched for H2AK119ub1 (peaks) in replicated samples are represented. (**F-I**) DNAme and H2AK119ub1 signal (log(ChIP/Input)) on *Xenopus laevis* genome CpG island (F) and repeat element (H). Z-score of H2AK119ub1 enrichment on CpG island (G) and repeat element (I) in sperm, replicated sperm and replicated USP21 sperm. *** p<0.005, permutation test. Numbers of H2AK119ub1 peak overlapping the genomic feature are indicated for each cell type.

To then determine the functional relevance of sperm derived H2AK119ub1 to embryo development, we took advantage of the availability of paternally derived embryos in frog. As described previously, such embryos develop solely on chromatin derived from sperm and provide a direct measure of the paternal contribution to early development^9^. We generated paternally derived embryos from sperm pre-incubated in control or USP21 containing oocyte extracts and monitored their development (**Fig. 5A**)^31^. From gastrulation onwards, we observed a decrease in the developmental potential of USP21 treated-compared to control treated-sperm derived embryos (**Fig. 5B**). We conclude that besides its DNA content, sperm also contributes to the embryo an USP21 sensitive signal that is necessary for development. The observed defects occur around the time of zygotic gene activation suggesting that USP21 treatment associated developmental failure might relate to defective regulation of embryonic gene expression. To evaluate this possibility, we carried out transcriptome analysis of early gastrulae generated from pool of untreated, control extract-treated or USP21 extract-treated haploid embryos. We performed RNA-seq analysis using sperm, oocyte extract, and egg originating from different preparations for each of the biological replicates. Differential gene expression analysis indicates that the treatment of sperm with a control oocyte extract has minor effect on the resulting embryo transcriptome when compared to embryos generated from untreated sperm (**Fig. 5C**, left; **Suppl Fig. 10A-B**). By contrast, USP21 extract treatment leads to embryonic misregulation of 269 genes (**Fig. 5C**, right), the majority of which are upregulated in embryos derived from USP21 extract treated sperm compared to embryos derived from control extract treated sperm (74%, 200/269, **Fig. 5D, Suppl Fig. 10C, Suppl Table 4**). Thereafter, this set of genes found differentially expressed in embryos derived from sperm treated with USP21 oocyte will be referred to as USP21 sensitive genes. GO enrichment analysis highlight genes related to cell cycle processes in this gene set (**Suppl Table 4**). We next asked to which extend the expression pattern of USP21 sensitive genes identified on pooled embryos reflect the behaviour in single embryos. To that end, we carried out transcriptome analysis in single paternally derived gastrulae. We find that the expression pattern of USP21 sensitive gene observed in pooled embryos is replicated in half of the single embryos derived from USP21-treated sperm (4/8 USP21 treated sperm derived embryos, **Suppl Fig. 10D**). This heterogeneous transcriptome pattern matches the observed developmental outcome, with a fraction of gastrulae derived from USP21-treated sperm developing to the tadpole stage (**Fig. 5B)** and is reminiscent of the transcriptome heterogeneity detected in cloned embryos^32^. We then interrogated the developmental dynamic of USP21 sensitive genes. We observed a clear distinction between up- and down regulated USP21 sensitive genes (**Suppl Fig. 10E**). During development of embryos generated by *in vitro* fertilisation^33^, upregulated genes exhibit a strong maternal contribution followed by clearing prior to gastrulation (stage 10-12) and re-expression at tadpole stage (stage 25-40) while downregulated genes show increased expression around gastrulation. A recent study indicates that at ZGA, transcription occurs on genes with (maternal-zygotic, “MZ genes”) or without (zygotic, “Z genes”) a maternal contribution of transcripts^34,35^. Interestingly, we observed that USP21 sensitive genes are enriched for MZ genes, indicating that these genes exhibit zygotic transcription in the context of a maternally derived transcripts (**Suppl Fig. 10E-F)**. Taken together, these analyses support the hypothesis whereby USP21 mediated erasure of the repressive histone modifications H2AK119ub1 from sperm leads to precocious embryonic expression of a subset of genes that could in turn impact critical embryonic process such as cell division.

**Figure 5:**
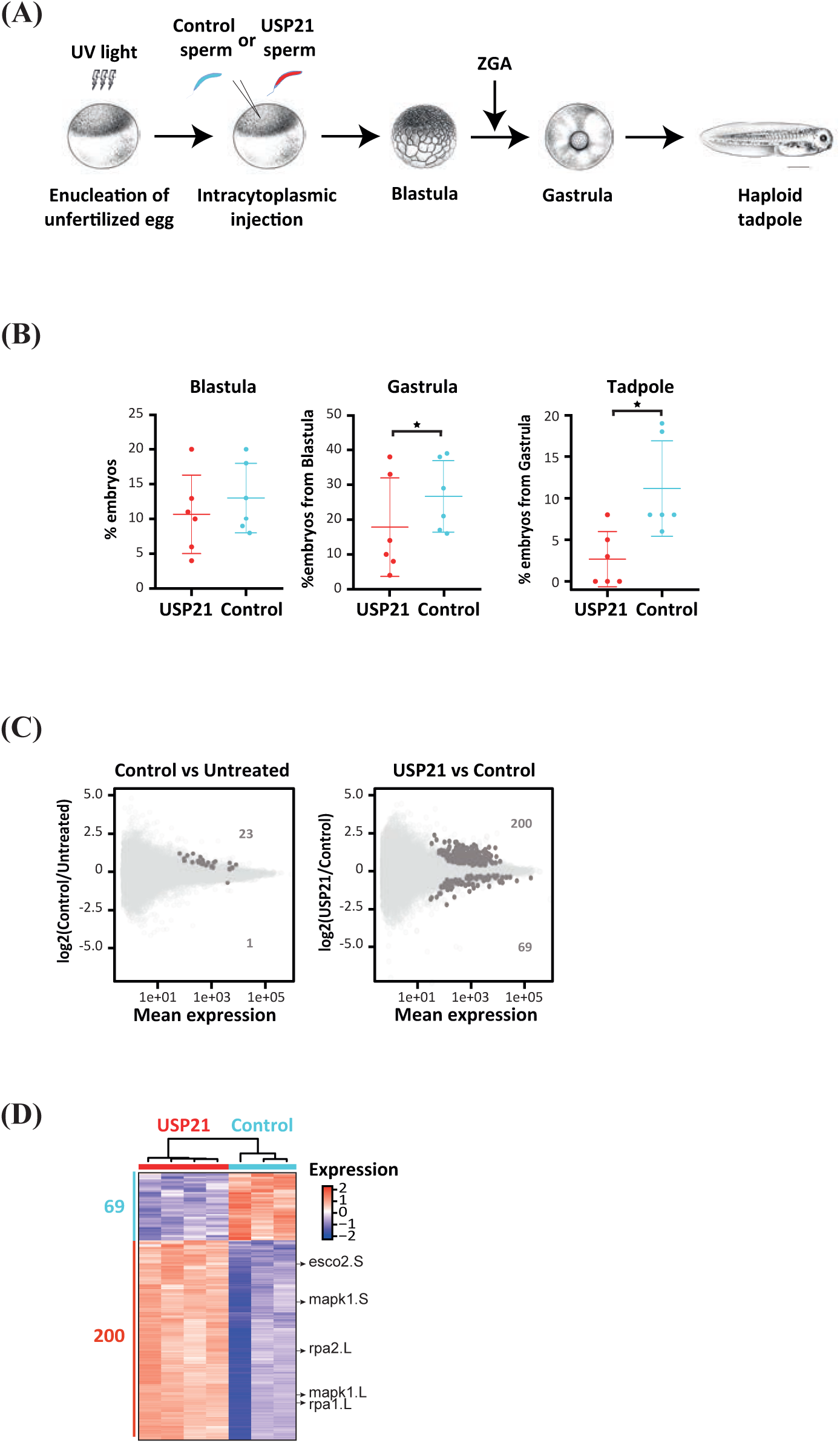
Sperm pre-treatment with a H2A deubiquitylase decreases developmental potential and leads to embryonic gene misregulation. (**A**) Experimental design for the generation of haploid embryos from sperm or USP21 treated sperm. *Xenopus* illustrations © Natalya Zahn (2022) from Xenbase (www.xenbase.org RRID:SCR_003280). (**B**) Developmental progression to blastula, gastrula or swimming tadpole stage of haploid embryos derived from a USP21 or control oocyte extract treated sperm. Chi-squared test, * p-value<0.05. Median and SD values are shown. (**C**) MA-Plot showing as y-axis the log foldchange (Log2FC) of gene expression between control oocyte extract treated sperm *versus* untreated sperm (left) and between USP21 oocyte extract treated sperm derived embryo and control oocyte extract treated sperm (right). x-axis is the averaged gene expression across samples.

To evaluate this possibility further, we investigated to which extent H2AK119ub1 genomic distribution associate with embryonic gene sensitivity to sperm USP21 treatment. We observed an enrichment of sperm H2AK119ub1 peaks on USP21 sensitive gene body (**Suppl Fig. 10G**). Additionally, using a genome partitioning approach (hidden Markov model) we also observe that USP21 sensitive genes are associated with genomic regions exhibiting H2AK119ub1 either in sperm or in sperm following exposure to egg extract (**Suppl Fig. 10H).** However, many more genes harbour this modification in sperm, without being affected by USP21 treatment. We therefore next interrogated the association of USP21 sensitive genes with different types of H2AK119ub1 peaks stratified according to association with other histone PTMs, namely H3K4me3 and H3K27me3 (as in **Fig. 2E-F**). We detect an association of USP21 sensitive gene features with either H2AK119ub1 only peaks (H2AK119ub1+; H3K4me3-; H3K27me3-) or trivalent H2AK119ub1 peaks (H2AK119ub1+; H3K4me3+; H3K27me3+) (**Fig. 6A**). Strikingly however, USP21 sensitive genes show a strong enrichment for bivalent H2AK119ub1/H3K4me3 and depletion for bivalent H2AK119ub1/H3K27me3 chromatin configuration. This analysis suggests that the effect on embryonic gene expression of sperm USP21 treatment arise from erasure of H2AK119ub1 in the context of a bivalent H2AK119ub1/H3K4me3 chromatin status. We also observed that, post-replication, H2AK119ub1 and H3K4me3 signals around USP21 sensitive gene TSSs are higher than around the bulk of *X. laevis* genes TSSs (**Fig. 6B**). Importantly, the significant enrichment of USP21 sensitive genes promoter and gene body for a bivalent H2AK119ub1/H3K4me3 chromatin configuration reported in sperm holds true after sperm replication (**Suppl Fig. 10I**). This association of USP21 sensitive genes with H2AK119ub1 is also observed in embryos at ZGA, with such trend still present but less pronounced later during embryogenesis (**Suppl Fig. 6D**). We then took advantage of a published dataset describing the consequences of expressing the H2A deubiquitylase BAP1 in early mouse embryos to investigate if such a relationship is conserved across specie^19^. Using a hidden Markov model, we indeed found that gene identified as BAP1 sensitive in mouse two cell embryos are enriched in genome domains exhibiting H2AK119ub1 marking in sperm (**Suppl Fig. 10J-K**). Altogether these observations support a role for sperm derived H2AK119ub1 in the regulation of the expression of a subset of embryonic genes.

**Figure 6:**
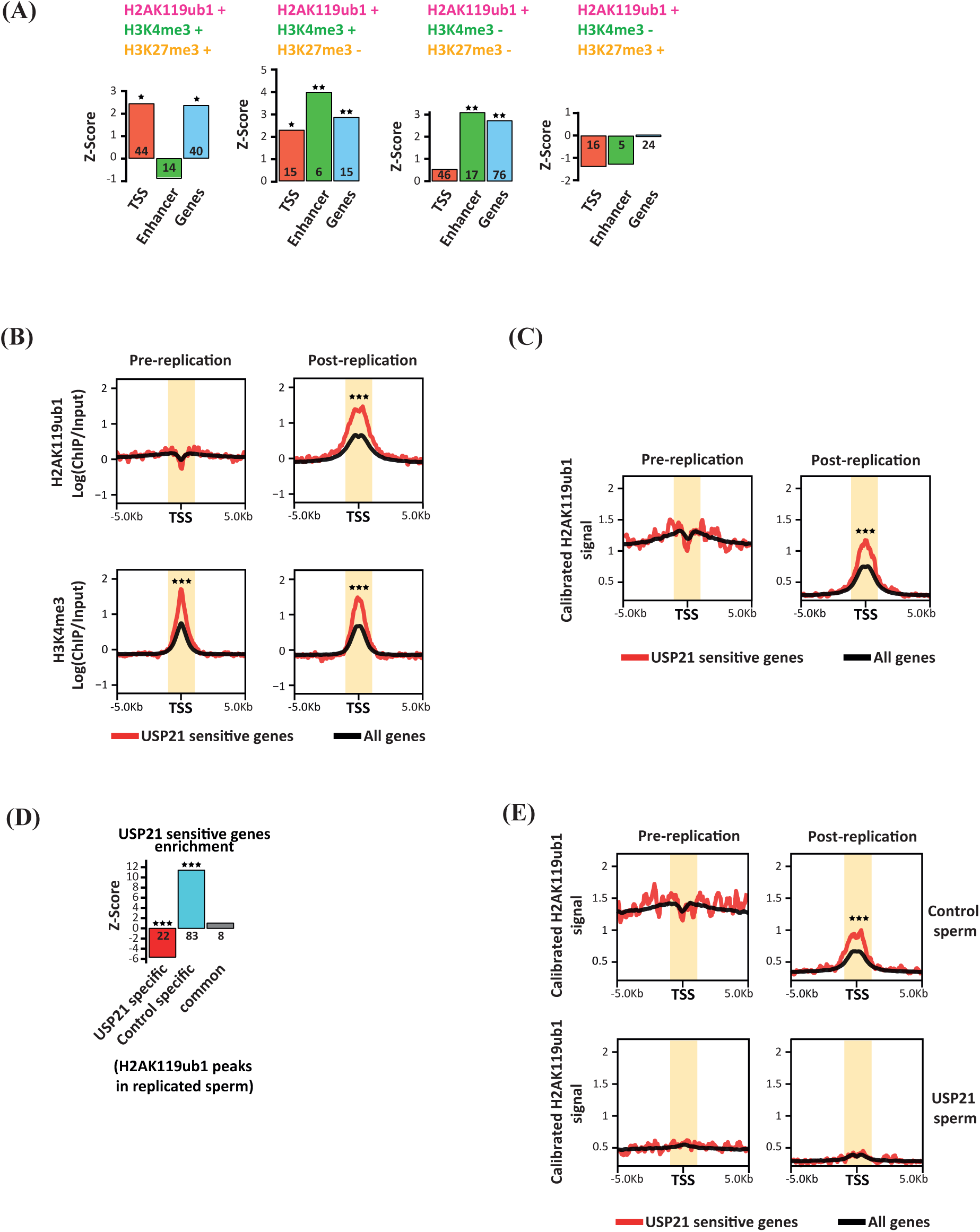
Genes sensitive to sperm deubiquitylase treatment are H2AK119ub1 target. **(A)** Heatmap showing expression of differentially expressed gene in gastrulae derived from USP21 oocyte extract treated sperm *versus* that derived from control oocyte extract treated sperm. **(B)** Z-score of different H2AK119ub1 peaks categories enrichment on TSS, enhancer, and gene body of USP21 sensitive genes. * p-value<0.05; ** p-value < 0.01, permutation test. Number of H2AK119ub1 peak overlapping the USP21 sensitive gene feature are indicated for each category. **(C)** H2AK119ub1 and H3K4me3 signal on TSS +/- 5kB for all genes (black), USP21 sensitive genes (red) before and after sperm replication. *** p-value<0.001, Wilcoxon-Mann-Whitney test on TSS+/1kb interval (shaded orange area). **(D)** Calibrated H2AK119ub1 ChIP-seq signal on TSS +/- 5kB for all genes (black), USP21 sensitive genes (red) before and after sperm replication. *** p-value<0.001, Wilcoxon-Mann-Whitney test. **(E)** Z-score of USP21 sensitive gene enrichment on H2AK119ub1 peaks gained (USP21 specific), lost (Control specific) or maintained (common) in replicated USP21 treated sperm *versus* replicated control sperm. ***p-value<0.001, permutation test. Number of USP21 sensitive gene overlapping each category is shown. **(F)** Calibrated H2AK119ub1 ChIP-seq signal on TSS +/- 5kB for all genes (black), USP21 sensitive genes (red) before and after replication using a control oocyte extract treated sperm or USP21 treated sperm. *** p-value<0.001, Wilcoxon-Mann-Whitney test.

We next considered the possibility that while a local accumulation of H2AK119ub1 could be maintained post-replication, sperm modified histones could have undergone replicative dilution. In other word, during replication the local accumulation of histone along the genome could be maintained albeit at a reduced level, for example through redistribution of sperm derived modified histones between the replicated strands of DNA. Mechanistically, this is important as in frog ∼8 rounds of replication separate fertilisation form the main wave of ZGA and could therefore leads to complete loss of sperm derived modified histones. Therefore, to compare H2AK119ub1 level pre- and post- replication we calibrated the signal *via* semi-synthetic H2AK119ub1 nucleosome spiked-in used in the ChIP-seq (see M&M) (**Fig.6C**). We observed a global decrease of H2AK119ub1 signal on TSSs upon egg extract treatment, suggesting that sperm chromatin replication induces redistribution of this modified histones at these locations (black line, **Fig. 6C**). When considering USP21 sensitive gene however, we observed a different trend. While an overall decrease of H2AK119ub1 signal is also detected upon replication, H2AK119ub1 loss is clearly less pronounced on the TSSs of this subset of gene than seen genome wide (red line, **Fig. 6C),** a feature shared with the broader set of gene that have maternal-zygotic contribution (green line, **Suppl Fig. 10L**). These analyses indicate that USP21 sensitive genes are indeed associated with a sperm H2AK119ub1 configuration that persists through egg factor mediated remodelling of the paternal epigenome. Lastly, we asked how sperm USP21 treatment induced disruption of paternal epigenome remodelling by egg factors affects USP21 sensitive genes. We found that the shift of H2AK119ub1 away from CpG towards repeat elements induced by USP21 treatment (as in **Fig. 4E**) significantly deplete post replication association of USP21 sensitive genes from this histone modification (**Fig. 6D**). Analysis of gene TSSs also showed that the preferential post replication retention of H2AK119ub1 on USP21 sensitive genes is lost in response to H2AK119ub1 sperm editing (**Fig. 6E**).

Taken together our results reveal that the setup of paternally derived zygotic chromatin is dictated by component of the sperm epigenome and that this process is required for proper expression of a subset of embryonic genes.

## DISCUSSION

In this work, we found that in *Xenopus laevis*, paternal H2AK119ub1 genomic distribution is extensively remodelled upon exposure to egg factors, reminiscent of the situation found in the mouse^19^. In this dynamic epigenetic landscape, a subset of genomic location is nevertheless found to maintain H2AK119ub1 following sperm replication in extract as well as following early embryonic division. This suggests that the mechanism recently described in cultured cells and demonstrating recycling of H2AK119ub1 between daughter cells^67^ could apply to the transmission of H2A associated epigenetic cue from the gametes to the dividing cell of the frog embryos. Besides those stable sites of H2AK119ub1 accumulation, sperm incubation in egg extract also triggers H2AK119ub1 deposition on new genomic sites. Whether this remodelling requires replication is not known since sperm treatment with egg extract triggers many processes besides replication. Once established those new sites appear stable through replication as they could be detected in late cleavage stage embryos. However, our study is limited by the lack of knowledge about H2AK119ub1 genomic distribution during xenopus laevis pre-ZGA developmental period. Particularly, we are lacking in vivo data describing the H2AK119ub1 landscape immediately after fertilisation and during the first round of embryonic divisions. Therefore, mechanistically we don’t know if the remodelled H2AK119ub1 distribution found after in vitro sperm replication, and in vivo after 8 round of cell division, is faithfully transmitted throughout cleavage stage or transiently adopt a different configuration during that period.

Egg factor dependent accumulation of H2AK119ub1 occurs around unmethylated CpG island and/or TF binding sites (*i.e.* USF1/2), genomic features that are known target of PRC1 activity in other systems^30^. Besides transcription factors, Kdm2b a component of vPRC1 is involved in more broadly targeting the H2AK119 ubiquitylation activity of this complex to unmethylated CpG island in ESC^30^. Kdm2b is a likely candidate for a role in the observed H2AK119ub1 remodelling following exposure to egg components as it is maternally provided as protein in frog (https://www.xenbase.org/xenbase/gene/geneExpressionChart.do?method=dr <u>awProtein</u>). In mouse embryonic stem cell, the establishment of polycomb repressive domain occurs in a stepwise manner with a variant PRC1 complex initiating their formation through ubiquitylation of H2AK119, followed by recruitment of the PRC2 complex and subsequent methylation of H3K27me3. Such sequential events have also been described during Neural Progenitor Cell differentiation^36^ and X chromosome inactivation^37^. Recently, this sequence of events have been investigated in the mouse and zebrafish embryos^19,20,38^. In these embryos H2AK119ub1 is also required for the deposition of H3K27me3 and transcriptional repression around the time of zygotic gene activation. Our observation that H2AK119ub1 is present in pre-ZGA embryo in the absence of H3K27me3 argues for the existence of a similar mechanism in frog^39,40^.

Interference with H2AK119ub1 in the mouse, either by knocking down PRC1 component^38^ in oocyte, or erasing H2AK119ub1 from embryos^19,20^ leads to mis-regulation of genes at ZGA and is associated with embryonic lethality. While clearly demonstrating a role for H2AK119ub1 in regulation of transcription around ZGA, these approaches entail prolonged epigenetic interference during either gametogenesis or early embryogenesis and, as a consequence, do not enable to determine the extent to which embryonic H2AK119ub1 is instructed/inherited from sperm and/or oocyte H2AK119ub1^9,11,12,19,20,38^. Here, by editing the sperm epigenome, after completion of spermiogenesis and prior to exposure to egg factors, we circumvent these limitations. We demonstrate that sperm contribute to the embryo a USP21 sensitive cue required for paternal H2AK119ub1 remodelling by egg factor, embryonic gene expression, and developmental competence (**Fig. 7**). We find that egg factor mediated accumulation on paternal CpG island depends on H2AK119ub1 delivered by the sperm. Strikingly, we observe that erasure of H2AK119ub1 from sperm leads to a global shift in genomic accumulation from CpG islands to repeat elements. Given the propensity of vPRC1 to be targeted to and to spread from existing H2AK119ub1 location^41^ it raises the possibility that pre-existing sperm derived “docking sites” could limit chromatin sampling by the polycomb complex and prevent H2AK119ub1 deposition at genomic location where targeting is usually limited such as the repeat elements. Embryos derived from H2AK119ub1 edited sperm exhibit transcriptional misregulation, mostly upregulation, compatible with the repressive function of this histone PTM on transcription. A similar observation is observed in mouse when H2AK119ub1 deubiquitylase is expressed at fertilization^19^, or in zebrafish when Rnf2 is chemically inhibited^20^. Given the extent of egg factor mediated H2AK119ub1 accumulation on gene promoters observed in frog, mouse or zebrafish, it is somewhat surprising that only a limited number of genes are mis-regulated when its distribution is massively altered. We identify an association of misregulated genes with H2AK119ub1 in frog sperm and find that this association is preferentially maintained on these genes upon sperm chromatin replication, further supporting a direct role for this histone post-translational modification in the epigenetic programming of sperm for regulation of embryonic transcription. We cannot rule out that a part of the genes is indirectly affected, for example as downstream target of a “primary” H2AK119ub1 target. Alternatively, we provide evidence that gene sensitivity to H2AK119ub1 erasure from sperm occurs when genes are associated with H3K4me3, a histone PTM known to act as a vector of intergenerational information^9–11,42,43^. It is likely that additional epigenetic features besides H3K4me3 contribute to gene sensitivity to H2AK119ub1 erasure from sperm. Additionally, we observe that most affected genes are normally expressed late in embryogenesis (tadpole stage) a feature also seen in mouse with a large proportion of gene sensitive to H2AK119ub1 depletion expressed post-implantation^8^. It is therefore possible that the effect of H2AK119ub1 expression is only partially revealed at early embryonic stage since transcription factors targeting these genes might not be present.

**Figure 7:**
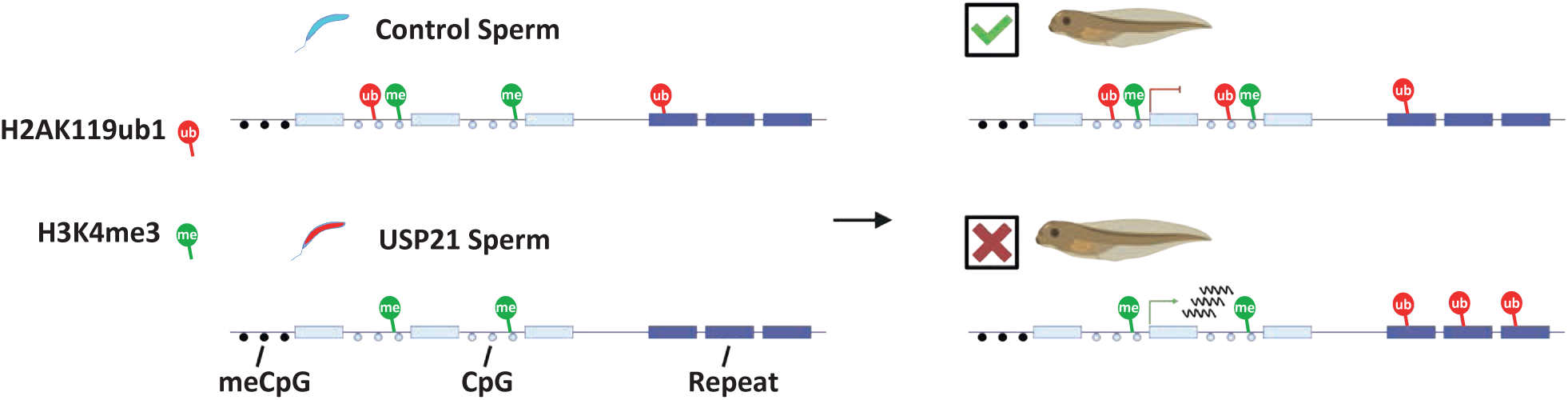
Model of H2AK119ub1 epigenetic programming of sperm for embryonic development in Xenopus laevis.

Our work advance knowledge form previous studies by showing that part of H2AK119ub1 found in embryos is paternally instructed. This necessary function of sperm derived modified histones has important bearing on our understanding of inter/trans generational transmission of epigenetic information. One could indeedenvisage that such a naturally occurring mechanism of epigenetic information transmission to the embryos could be hijacked to transmit environmentally induced sperm epigenetic cues. Based on the very dynamic nature of histone PTMs around fertilisation, it is likely that transmission to the embryos will be context- (combination of marks) and location- dependent. Future studies will aim at evaluating to which extent sperm histones PTMs differ between individuals and to which extent they are maintained in embryos.

## FIGURE LEGENDS

**<u>Supplemental Figure 1:</u> *Quantification of Western blot data*.**

**(A)** Ratio of fluorescence intensity detected by WB for H2AK119ub1/H2A (left) H2AK119ub1/H4 (right) and H2A/H4 (middle). The same data as Fig.1G are plotted with the addition of a solid line linking paired sample quantified in the same WB experiment. **(B)** H2AK119ub1 fluorescence intensity detected by WB in untreated, control oocyte extract treated sperm, USP21 extract treated sperm. The same data as Fig.2E are plotted with the addition of a solid line linking paired sample quantified in the same WB experiment.

**<u>Supplemental Figure 2:</u> *Pooling strategy for ChIP-seq replicates***

**(A)** Genome browser view of the log(ChIP/Input) for H2AK119ub1 around the *hoxa.L* cluster on 3 sperm biological replicates. (**B**) Spearman correlation of sperm and replicated sperm H2AK119ub1, H3K27me3 and H3K4me3 ChIP-seq replicates. (**C**) (**D**) Heatmap illustrating the distribution of H2AK119ub1 and H3K4me3 in sperm and replicated sperm replicates and in the corresponding pooled sample obtained by subsampling. Log(ChIP/Input) signal is shown for TSS (C) and CpG (D), respectively. (**E**) Fingerprint plot of Input, H3K4me3, H3K27me3, and H2AK119ub1 sperm ChIP samples.

**<u>Supplemental Figure 3:</u> *Peak calling using RECOGNICER and SICER/MACS2*.**

**(A)** Overlap of peaks called using RECOGNICER, SICER, and MACS2. Numbers in red indicate RECOGNICER specific peaks *versus* those overlapping with those of SICER and/or MACS2. Diagram scaling is based on RECOGNICER peaks. Blue and green numbers correspond to the same comparison using SICER or MACS2 as the reference set of peaks, respectively. (**B**) Heatmap of H2AK119ub1, H3K27me3, H3K4me3 signal (log(ChIP/Input)) for RECOGNICER, SICER, and MACS2 set of peaks. (**C**) Comparison of Z-scores obtained from permutation test using set of peaks based on peaks set on peak calling using RECOGNICER (top row, corresponding to data presented in Fig. 2K, Fig. 2M, Fig. 3H and Suppl Fig. 10G) or SICER (bottom row). (**D**) Genomic distribution of sperm H2AK119ub1 peaks according to overlap with H3K4me3 and/or H3K27me3 peaks when calling peak using RECOGNICER (top, corresponding to data presented in Fig. 1I) or SICER (bottom) **(E)** Z-score indicating enrichment of different H2AK119ub1 peaks categories on TSS +/-5kb, enhancer, and gene body of USP21 sensitive genes. Top row is using RECOGNICER (already shown in Fig. 3E) while bottom row is using SICER for H2AK119ub1 peak calling.

**<u>Supplemental Figure 4:</u> *H2AK119ub1 is mostly associated with nucleosomes while H3K4me3/H3K27me3 are mostly associated with sub-nucleosomes in Xenopus laevis sperm.*** (**A**) Schematic representation of *Xenopus laevis* chromatin remodelling from nucleosomes to a mixture of nucleosomes and sub-nucleosomes during spermiogenesis. **(B)** Fragment size distribution obtained after paired-end sequencing of chromatin immunoprecipitated with antibodies against H2AK119ub1 (pink) in sperm and spermatid, and against H3K4me3 (green) and H3K27me3 (yellow) in sperm. Matching Input fraction are shown in grey.

**<u>Supplemental Figure 5:</u> *Sperm H2AK119ub1 fate following exposure to egg factors***

**(A)** Heatmap of H2AK119ub1 (pink), H3K4me3 (green), and H3K27me3 (yellow) log(ChIP/Input) signal ratio on genes TSS from spermatid, sperm prior and after egg extract treatment, and gastrula stage 12 embryo. (**B**) Scaled Venn diagram of H2AK119ub1, H3K4me3 or overlapping H2AK119ub1/H3K4me3 peaks in spermatid, sperm, and sperm exposed to egg factors. Percentage of peaks that are maintained at each developmental transition are indicated underneath. (**C**) Top: boxplot showing calibrated H2AK119ub1 on H2AK119ub1/H3K4me3 peaks lost (375), maintained (5222) or gained (10442) following exposure to egg factors (top). Bottom: peak size (blue dots) as well as H2A (red dots) in each peak category. **(D)** Genomic distribution of sperm peaks before or after replication, as well as the subset of peaks lost, maintained, or gained during replication.

**<u>Supplemental Figure 6:</u> *H2AK119ub1 distribution from spermiogenesis to late embryogenesis*.**

**(A)** Pearson correlation of H2AK119ub1 ChIP in spermatid, sperm, sperm replicated in egg extract, blastulae at ZGA, early gastrula, and late neurula. (**B**) (**C**) Heatmap of H2AK119ub1 log(ChIP/Input) signal for TSSs (B) and CpG islands (C) in spermatid, sperm, sperm replicated in egg extract, blastulae at ZGA, early gastrula, and late neurula. **(D)** Metaplot analysis of H2AK119ub1 log(ChIP/Input) signal for TSSs of all genes (black) or USP21 sensitive genes (red) in spermatid, sperm, sperm replicated in egg extract, blastulae at ZGA, early gastrula, and late neurula. *** p-value<0.001, Wilcoxon-Mann-Whitney test on TSS+/1kb interval (shaded orange area).

**<u>Supplemental Figure 7:</u> *Effect of oocyte extract treatment on sperm chromatin***

**(A)** Quantitation of H3K4me3, H3K9me3 and H3K27me3 to H4 WB signal ratio in untreated sperm, USP21 treated sperm or control oocyte extract treated sperm, **(B)** Capillary electrophoresis analysis of DNA fragments obtained following MNAse digestion of somatic cell (yellow), egg extract treated sperm (dark blue), sperm (blue), control oocyte extract treated sperm (cyan), and USP21 extract treated sperm (red). (**C**) Fragment size distribution obtained after paired-end sequencing of DNA fragments obtained following MNase digestion of sperm (top row), control oocyte extract treated sperm (middle row) or USP21 treated sperm (bottom row) without (left column) or with (right column) subsequent incubation in egg extract. (**D**)(**E**) Heatmap of H2AK119ub1 and H3K4me3 log(ChIP/Input) signal for TSSs (D) and CpG islands (E) in sperm and sperm treated with control oocyte extract.

**<u>Supplemental Figure 8:</u> *Fate of sperm H2AK119ub1 and H3K4me3 following exposure to egg factors*.**

**(A)** Experimental design for monitoring modified histones fate from control oocyte extract or USP21 treated sperm. **(B)** Heatmap of H2AK119ub1 (pink), H3K4me3 (green) and H3K27me3 (yellow) log(ChIP/Input) signal in spermatid, control oocyte extract- and USP21- treated sperm prior and after egg extract treatment, and gastrula stage 12 embryo. Heatmap includes the same genomic location as in Fig. 2B. **(C)** Heatmap of H2AK119ub1 and H3K4me3 signal log(ChIP/Input) in sperm after treatment with control oocyte extract or USP21 oocyte extract without or with subsequent replication in egg extract. All genomic locations enriched for H3K4me3 (peaks) in replicated samples are shown. (**D**) Transcription factor motifs enriched in peaks that are lost, maintained, or gained upon replication of control versus USP21 treated sperm (clusters from Figure 4E).

**<u>Supplemental Figure 9:</u> *H2AK119ub1 is enriched on repeat elements of the genome following exposure of USP21 treated sperm to egg factors.***

**(A)** Metaplot analysis of H2AK119ub1 log(ChIP/Input) signal on all CpG islands (10630 elements) and all repeat elements of the genome (313338 elements). * p- value<0.005, Wilcoxon-Mann-Whitney test is indicated with an asterisk in black (replicated control sperm *versus* replicated USP21 sperm), blue (replicated control *versus* control sperm) or red (replicated USP21 sperm *versus* control sperm). (**B**) same analysis as in (**A**) but splitting repeat elements according to subtypes.

***<u>Supplemental</u>* <u>Figure 10:</u> *Genes misexpressed in embryos upon H2AK119ub1 removal by deubiquitylases are associated with H2AK119ub1 in sperm.***

**(A)** Principal Component analysis of replicate RNA-seq samples used in this study (USP21, USP21 extract treated sperm derived gastrulae; Control, control oocyte extract treated sperm derived gastrulae; Untreated, untreated sperm derived gastrulae). **(B)** Hierarchical clustering of replicate samples used in this analysis. **(C)** Heatmap depicting expression of USP21 sensitive genes in replicates from USP21 extract treated sperm derived gastrulae, control extract treated sperm derived gastrulae, as well as gastrulae derived from untreated sperm. **(D)** Heatmap of gene expression in single haploid gastrula derived from control- or USP21-treated sperm. Expression is shown for the USP21 sensitive gene set identified on embryos pool (Fig. 3D). **(E)** USP21 sensitive gene set expression during early *Xenopus laevis* embryo development (data from^33^). Expression is shown for oocyte, egg, blastulae (stage 8-9), gastrulae (stage 10- 12), neurula (stage 15-20), as well as early to late tailbud (stage 25-40) are indicated. The right-hand side panel shows the expression of USP21 sensitive gene in gastrulae derived from control- and USP21-treated sperm. **(F)** Violin plot of gene expression in egg (log of normalised gene count) for all genes (black), USP21 sensitive genes (red), maternal genes (cyan), maternal and zygotic genes (green) and zygotic only gene (orange). USP21 sensitive genes are enriched for genes with maternal and zygotic expression in *Xenopus laevis*. The table below indicates p-Value resulting from a permutation test to evaluate enrichment of USP21 sensitive gene for either maternal-zygotic (MZ) or zygotic only (Z) genes. MZ and Z dataset taken from^34,35^. **(G)** Z-Score of H2AK119ub1 peak enrichment on TSS, enhancer, and gene body of USP21 sensitive genes. *p-value<0.05; permutation test. **(H)** Left, Heatmap representing H2AK119ub1 dynamics of each ChromHMM type in spermatid, sperm, replicated sperm, and gastrulae embryos. Color intensity represents H2AK119ub1 signal for a given state and cell type. Right, Z-score indicating enrichment for USP21 sensitive genes in genomic regions associated with the different chromHMM states. *p-value<0.05; **p-value < 0.01, ***p-value<0.001, permutation test. **(I)** Z-score of different H2AK119ub1 peaks sets enrichment on TSS, enhancer, and gene body of USP21 sensitive genes. Z-score are shown for sets corresponding to H2AK119ub1 peaks that are lost, maintained or gained following incubation of sperm in egg extract as well as for the subset of maintained H2AK119ub1 overlapping H3K4me3 peaks. *p-value<0.05; **p-value < 0.01, ***p-value<0.001, permutation test. **(J)** Experimental scheme used in Chen *et al.*, Nature genetics, 2021^19^. (**K**) Left, Heatmap representing H2AK119ub1 dynamics of each ChromHMM type in mouse embryos and gametes based on data generated^19^. Color intensity represents H2AK119ub1 percentage for a given state and cell type. Right, Z-score indicating enrichment for BAP1 sensitive genes identified in^19^ in genomic regions associated with the different chromHMM states. *p-value<0.05; **p- value < 0.01, ***p-value<0.001, permutation test. (**L**) Calibrated H2AK119ub1 ChIP signal on TSS +/- 5kb for all genes (black), USP21 sensitive genes (red), maternal genes (cyan), maternal zygotic genes (green) and zygotic genes (orange) before and after sperm replication. *** p-value<0.001, Wilcoxon-Mann-Whitney test.

**<u>Supplemental Table 1:</u> *Histone PTMs ChIP-seq***

**<u>Supplemental Table 2:</u> *Gene Ontology and TF motif analysis related to* Fig.2**

**<u>Supplemental Table 3:</u> *Gene Ontology related to Supplemental Fig.8***

**<u>Supplemental Table 4:</u> *Control- and epigenetically edited- sperm derived embryos RNA-seq data***

**<u>Supplemental Table 5:</u> *Repeat element analysis***

**<u>Supplemental Table 6:</u> *Extended Permutation test data***

## MATERIALS AND METHODS

### Contact for Reagent and Resource Sharing

Further information and requests for resources and reagents should be directed to and will be fulfilled by the Lead Contact, Dr J. Jullien.

### Experimental Model and Subject Details

Mature *Xenopus laevis* females were obtained from Tefor Paris-saclays (151 Rte de la Rotonde, 91400 Saclay, France, https://tefor.net/tefor-paris-saclay-nos-equipes/tefor-paris-saclay-aqua/). Our work with Xenopus complied with all relevant ethical regulations for animal testing and research. Work with frog was covered under the licence #202212131604754/APAFIS 39794V2 and frog husbandry and all experiments were performed according to the relevant regulatory standards. Animals were kept in a recirculating system (ALS, https://www.aquariumlocationservices.fr/professionnels-elevages-aquatique-aquariophilie/) at a density of one adult/3l, with ∼10% water change per day. Water was sequentially filtered with mechanical pad sump filter, nitrifying bacteria filter, and UV sterilized. Water quality parameters were as follow: temperature 17-22°C; PH 6-8. Photoperiod was set to 12h ON/12h OFF. Frogs are fed twice per week with NEO MARIN coul 4 - X4 - S20 (Le Gouessant, https://www.legouessant.com/). Unconsumed food was removed 30 min after the start of feeding.

### Oocyte extract

Fresh ovaries were manually dissociated in 15-25 oocytes clumps. 3-5 mL of dissociated ovary fragments were transferred into 12.5 mL 1X Modified Barth’s Saline (MBS, 88 mM NaCl, 1 mM KCl, 10 mM Hepes, 2.4 mM NaHCO3, 1.7 mM MgSO4, 1.3 mM Ca(NO3)2, 0.8 mM CaCl2) supplemented with 250 µL of 5 mg/mL Liberase (Roche, 05 401 127 001) and incubated for 2h30-3h at room temperature (RT) with gentle rocking. Defolliculated oocytes were washed three times with 1X MBS and healthy stage 5-6 oocytes manually sorted. Oocytes were then injected in the vegetal pole with 25 ng of USP21-NLS-HA mRNA, or control GFP mRNA and incubated for 3 days at 18°C in 1X MBS, renewing medium and discarding dead cells every day. ∼500-1000 oocytes were collected in an Eppendorf tube on ice, the excess 1X MBS discarded and replaced with 1X MBS supplemented with Protease Inhibitor Cocktail (Sigma, P2714-1BTL). The oocytes were then stacked up by centrifugation at 100 g for 1 min at RT and excess medium carefully discarded. Oocytes were then thoroughly vortexed and centrifuged at 21 000g for 10 min at 4°C. The clear intermediate phase between yolk (top) and pigment (bottom) was retrieved. The collected fraction was further cleared by an additional centrifugation at 21 000 g for 10 min at 4°C. The oocyte extract was then aliquoted and stored at −80°C.

### Oocyte extract treatment

Permeabilized sperm stored in SSB at −80°C were thawed, washed with two volumes of ice cold 1X PBS and centrifuged at 800 g for 8 min at 4°C. 2M sperm in 1X PBS were centrifuged at 21000 g for 1 min at 4°C. The supernatant was discarded, and the pelleted sperm resuspended in 10 µL of oocyte extract on ice. The sperm incubated in the oocyte extract for 20-40 min at 23°C on a heating block. Treatment was stopped by addition of 90 µL of ice cold 1X PBS and centrifugation at 21000 g for 1 min at 4°C. The supernatant discarded and the remaining pellets resuspended in SSB for −80°C long term storage.

### Egg extract treatment

Interphasic extracts were generated as previously described^45^ .For the quantification of DNA replication in individual nuclei in egg extract, 33 μM rhodamine dUTP (Roche, 11534378910) was added at the indicated times (20, 40, 60, 80 mins after incubation) for 5 min as indicated, and the reaction was stopped with 1X PBS and fixed in 4% paraformaldehyde. The reaction was spun through a 30 % sucrose cushion in PBS. The sperm cells were rinsed twice in 1X PBS for 10 min and mounted in Vectashield mounting medium with DAPI (Vector laboratories, H-1200). Counting of rhodamine positive cells were performed on an epifluorescence microscope.

### Protein extraction and Western Blot analysis

Sperm samples stored in SSB were washed in 1X PBS and 1M sperm pellets were resuspended in 30 µL 2X Blue Eye (600 mM NaCl, 1 % Triton X100, 0.5 % Na-Deoxycholate, 4 % Sodium Dodecyl Sulfate, 20 % Glycerol, 0.0004 % Bromophenol blue, 175 mM Tris-HCl, 10 % Betamercapto ethanol), boiled at 96°C for 5 min and centrifuged at 21000g for 5 min at 4°C. The supernatant was transferred to a new tube and directly use for analysis or stored at −20°C. 4-20 % mini-PROTEAN® TGX (Biorad, #4561094) were loaded with 10 µL of sample and western blot analysis were performed according to standard protocols. For blotting nitrocellulose membranes, a semi-dry transfer was used (Biorad, Trans-Blot® SD Semi-Dry Transfer Cell) 30-40 min at 25V. The following primary antibodies were used: anti-H2A (1:2500, Cell Signalling, 3636), anti-H2AK119ub1 (1:2500, Cell Signalling, 8240), anti-H3K4me3 (1:2500, Abcam, ab8580), anti H3K9me3 (1:2500, Abcam, ab8898), anti-H3K27me3(1:2500, Cell Signalling, 9735S), anti-H4 (1:2500, Abcam, ab31830) and anti-HA (1:2500, Sigma, H9658). Membranes were incubated with primary antibodies 2 h at RT or overnight (ON) at 4 °C. Secondary antibody are anti-rabbit (1:25000, ThermoFisher, A21076), anti-mouse IgG Alexa Fluor 680 (1:25000, ThermoFisher, A21058), Goat anti-rabbit Alexa 800 (1:25,000, ThermoFisher, A32735), or IRDye 800CW Goat Anti-Mouse IgG (H+L) (LI-COR, 926-32210) were used. The membranes were imaged either on a ChemiDoc MP Imaging System (Biorad) or an Odyssey scanner (LI-COR).

### Sperm Chromatin Immunoprecipitation (ChIP)

Chromatin fractionation and chromatin immunoprecipitation (ChIP) were performed as described before^10,44^ with slight modifications. 80 μL of Magnetic beads (M280 Sheep anti-rabbit IgG, Invitrogen, #11204D) were used per reaction and all wash steps were carried out with a magnet. Beads were washed with 1 mL of TE pH 8.0, lysis buffer (LB)(1:1 mixture of Buffer 1 and MNase buffer, see below), and incubated in 1 mL of LB with 100 μL of 10 mg/mL BSA at 4°C for 30 min (“pre-blocking”). Pre-blocked beads were washed with 1 mL of LB twice and resuspend in 80 μL of LB. Half of beads were incubated with antibody overnight and leftovers were stored in 4°C for “pre-clear chromatin” step. For ChIP, the following antibodies were used: anti-H3K4me3 (Abcam, ab8580), anti-H2AK119ub1 (Cell Signaling, 8240). One million digitonin permeabilised sperm nuclei or egg extract treated sperm nuclei were resuspended in 50 μL of Buffer 1 (0.3 M Sucrose, 15 mM Tris pH 7.5, 60 mM KCl, 15 mM NaCl, 5 mM MgCl2, 0.1 mM EGTA, 0.5 mM DTT) containing 0.125 ng H2AK119ub biotinyltated nucleosome (Epicypher #16-0020), followed by addition of 50 μL of Buffer 1 with detergent (Buffer 1 including 0.5 % NP40 and 1 % NaDOC). Samples were incubated for 10 min on ice. 100 μL of MNase buffer (0.3 M Sucrose, 85 mM Tris, 3 mM MgCl_2_, 2 mM CaCl_2_, 2.5 U of micrococcal nuclease: Roche 10107921001) was added in to each tube (0.5 million of cells per tube). Tubes were incubated at 37°C for 30 min in pre-warmed water bath. Reaction was stopped by adding 2 μL of 0.5 M EDTA pH 8.0 in the same order as started. Tubes were vortexed and placed on ice for at least 5 min. Supernatant and pellet were separated by centrifugation at 13000 rpm for 10 min at room temperature. Beads (stored at 4C for pre-clear chromatin) were washed with 1 mL of LB twice. Supernatant and EDTA-free Protease Inhibitor Cocktail were added into the beads and rotating on the wheel at 4°C for 60 min. 10% of volume of pre-cleared chromatin was taken as “Input” and stored at 4°C. Antibody conjugated beads were washed with 1 mL of LB twice. Pre-cleared chromatin was added to washed beads and incubated at 4°C for 6 h. After incubation, beads were washed with 1 mL of Washing buffer A (50 mM Tris-HCl pH 7.5, 10 mM EDTA and 75 mM NaCl) on the wheel at 4C for 5 min and buffer was discarded carefully. Subsequently, beads were washed wish 1 ml of Washing buffer B (50 mM Tris-HCl pH 7.5, 10 mM EDTA and 125 mM NaCl) and buffer was discarded carefully. Another 1 mL of Washing buffer B was added and transferred to 1.5 mL Protein lo-bind tubes (Eppendorf, 0030108116) with beads. Buffer was discarded carefully. 150 μL of elution buffer (1:9 mixture of 10 % SDS and TE) was added. Tubes were placed at 25°C on the Eppendorf thermomixer and shaken at 800 rpm for 15 min. Supernatant were taken and transferred to the new tube using magnetic stand. Another 150 μL of elution buffer was added and repeated this step (in total 300 µl). Input samples were diluted to 300 μL total by adding TE. 2 μL of 2 mg/ml RNase A (heat-inactivated or DNase-free) was added and incubated at 37°C for 30 min. 4 μL of 10 mg/ml Proteinase K was added and incubated at 55°C overnight. After incubation, tubes were shortly spun. 300 μL of Phenol/chloroform were added and vortexed for 30 sec. Samples were centrifuged at 13000 rpm for 10 min at room temperature. 1 µL of Glycogen, 1/10 volume of NaAcetate and 2 volumes of 100% EtOH were added and vortexed. Tubes were stored on dry ice for 20-30 min. Once frozen, tubes were centrifuged at 13000 rpm for 30 min at 4°C. Supernatant was discarded carefully and 500 μL of 70% EtOH was added. Tubes were centrifuged at 13000 rpm for 10 min at 4°C. Pellets were dried and suspended in 30 µL of ddH_2_O.

### Embryo Chromatin Immunoprecipitation (ChIP)

Batches of 200 in vitro fertilized embryos were fixed in 15 ml 1 % Paraformaldehyde in 1X MMR for 20min at room temperature followed by 4 washes with 1X MMR. Embryos were split in two and equilibrated at 4°C overnight in 500 µL HEG solution (50 mM HEPES-KOH pH 7.5, 1 mM EDTA, 20 % Glycerol), then excess buffer was discarded, and samples were transferred to −80°C. For chromatin extraction embryos were resuspended in 700 µL of sonication buffer (20 mM Tris-HCl pH 8.0, 70 mM KCl, 1 mM EDTA pH 8.0, 10% Glycerol, 5 mM DTT, 0.125% NP40, 1X complete protease inhibitors) and homogenized by pipetting up and down. 200 µL embryo extract were distributed in 1.5 mL tubes and sonicated in two sets of 15 min cycles (with 30 s on/off cycles). Sonicated samples were centrifuged at 12000 rpm for 5 min at 4 °C a Supernatants were retrieved and the centrifugation was repeated. Before ChIP, 180 µL (2x 10%) of the chromatin extract was taken as an Input control and stored at −20°C. For each ChIP, 720 µL of chromatin extract was mixed with 2 µL of anti-H2AK119ub1 antibodies (Cell Signalling, 8240) and incubated overnight on a rotating wheel. The next day, 40 µL (20 µL per IP) of Magnetic beads (M280 Sheep anti-rabbit IgG, Invitrogen, #11204D) were washed with 1X PBS using a magnet and incubated with 1mL of 1X PBS containing 1 mg/mL BSA for 2 h at 4°C on a rotating wheel. Blocking solution was discarded from the beads using a magnet and beads were resuspended in 200 µL (100 µl per IP) of incubation buffer (50 mM Tris-HCl pH 8, 100 mM NaCl, 2 mM EDTA, 1% NP40, 1 mM DTT, 1X Protease inhibitor). 100 µL of incubation buffer containing beads is transferred to chromatin/antibody mix and incubated for 4 h at 4°C. Samples were washed with Buffer 1 (50 mM Tris-HCl pH 8, 100 mM NaCl, 2 mM EDTA, 1 mM DTT, 1 % NP40, 0.1 % sodium deoxycholate, 1X complete protease inhibitors) for 5 min at 4°C on rotating wheel. Using a magnet, supernatant was discarded and beads subsequently washed with Buffer 2 (50 mM Tris-HCl pH 8, 500 mM NaCl, 2 mM EDTA, 1 mM DTT, 1 % NP40, 0.1 % sodium deoxycholate, 1X complete protease inhibitors), Buffer 3 (50 mM Tris-HCl pH 8, 100 mM NaCl, 250 mM LiCl, 2 mM EDTA, 1 mM DTT, 1 % NP40, 0.1 % sodium deoxycholate, 1X complete protease inhibitors), Buffer 1 after which immunoprecipitated chromatin was eluted using 400 µL (310 µL for Input fraction) of Elution buffer (0.1 M NaHCO3, 1 % SDS) overnight at 65°C. The next day, 4 µL of 20 mg/mL RNAse A was added to the chromatin solution and incubated for 2 h at 37°C. After incubation, tubes were shortly spun. 300 μL of Phenol/chloroform were added and vortexed for 30 sec. Samples were centrifuged at 13,000 rpm for 10 min at room temperature. 1 ul of Glycogen, 1/10 volume of NaAcetate and 2 volumes of 100% EtOH were added and vortexed. Tubes were stored on dry ice for 20-30 min. Once frozen, tubes were centrifuged at 13,000 rpm for 30 min at 4C. Supernatant was discarded carefully and 500 μL of 70% EtOH was added. Tubes were centrifuged at 13,000 rpm for 10 min at 4C. Pellets were dried and suspended in 30 *μ*l of ddH2O.

### *Xenopus* sperm collection *Xenopus* sperm collection

*Xenopus* sperm collection was performed as described before^10,44^. For each round of sperm purification, testes from six adult *Xenopus laevis* males were isolated and manually cleaned from blood vessels and fat bodies in 1X Marc’s Modified Ringers (MMR, 100 mM NaCl, 2 mM KCl, 1 mM MgSO_4_, 2 mM CaCl_2_, 3 mM HEPES, pH 7.4) using forceps and paper tissues. Subsequently, testes were torn into small pieces with forceps and homogenized with 2– 3 strokes of a Dounce homogenizer (one testis at a time). The cell suspension was then filtered to remove tissue debris and cell clumps using a 60 µm Mesh (CellTrics, cat. 04-0042-2317) and spun down at 800 rcf, 4°C, for 20 min. Supernatant was discarded and the cell pellet was resuspended in 12 mL of 1 × MMR. If any red blood cells were visible at the bottom of the pellet (as a result of incomplete removal of blood vessels), only the uncontaminated part of the pellet was recovered, taking extreme care not to disturb the red blood cells. Subsequently, step gradients of iodixanol (Optiprep; Sigma, D1556; 60% iodixanol in water) in 1X MMR final were manually prepared in pre-chilled 14 mL ultra-clear centrifuge tubes (Beckman Coulter, #344060) in the following order from the bottom to the top of the tube: 4 mL of 30 % iodixanol, 1 mL of 20 % iodixanol, 5 mL of 12 % iodixanol (all in 1 × MMR), and 2 mL of cell suspension in 1X MMR on top. Gradients were spun down in a pre-chilled SW40Ti rotor at 7500 rpm (10,000g), 4°C, for 15 min, deceleration without brake (Beckman Coulter Ultracentrifuge, Optima L-100XP). The pelleted fraction, containing mature sperm, was collected. Collected fractions were diluted six times with 1X MMR and collected by spinning first at 805 rcf, 4°C, for 20 min and then at 3220 rcf, 4°C, for 20 min to pellet remaining cells. Pelleted cells were subsequently permeabilized with Digitonin (10 mg/mL as a final concentration, Sigma, D141) for 5 min at room temperature as detailed before (Smith et al., 2006).

### Library preparation and sequencing

ChIP samples were subjected to ChIP-seq library preparation using QIAseq Ultralow Input Lib UDI-A Kit (Qiagen, 180497) and Agencourt AMPure XP beads (Beckman coulter, A63881). Samples pre- and post-library protocol were analyzed on a LabChip GX (PerkinElmer) using DNA 1KHS and 1K Assay LabChIP, respectively. Libraries were quantitated using NEBNext Library Quant Kit (NEB, E7630,). For RNA sequencing, RNAs were purified using Qiagen RNeasy Mini Kit (#74004) and used to generate cDNA sequencing libraries using an Illumina TruSeq Kit (#RS-122-2001), according to the manufacturer’s specifications. Paired-end sequencing was carried out either on a NOVA-seq 6000 or HiSeq 1500 sequencer (Illumina).

### RNA-Seq analysis

Adapters were trimmed and low-quality reads filtered using cutadapt (v1.18)^46^. Sample quality was confirmed with fastqc (v0.11.8)^47^. fastq were aligned to *Xenopus laevis* genome (v9.2) using STAR aligner (v2.7.3.a)^48^. Count table was generated using HTSeq-count (v0.12.3)^49^ and exon as genes reference. MultiQC (v1.9)^50^ summarized data pre-processing. Differential gene expression analysis of the count table was performed on R (v3.6)^51^ using DESeq2tools^52^. Details of the pipeline used can be found at https://gitlab.univ-nantes.fr/E114424Z/BulkRNAseq. Gene ontology analysis were carried out with gprofiler^53^ using human orthologs of *Xenopus laevis* genes recovered from Xenbase (https://www.xenbase.org/xenbase/static-xenbase/ftpDatafiles.jsp)^54^. *Xenopus laevis* development RNA-seq data were from published dataset GSE73430.

### ChIP-Seq analysis

Adapters were trimmed and low-quality reads filtered using cutadapt (v1.18)^46^. Sample quality was confirmed with fastqc (v0.11.9)^47^. Alignment was performed with bwa mem^55^ against *Xenopus laevis* (v9.2) or against a *Xenopus laevis* (v9.2) −601DNA hybrid genome generated with bwa index (v0.7.17-r1188). Reads extraction and analysis was performed with samtools view and samtools flagstat (v1.12)^56^. ChIP-Seq sample quality was assessed by Deeptools (v3.5.1)^57^ using plotFingerprint function and bamPEFragmentSize. Replicates quality was visualised with plotCorrelation using spearman correlation after genome fragmentation into bins of 10Kb. PCR duplicates were marked by Picard tools using MarkDuplicates command (https://broadinstitute.github.io/picard) and then removed with samtools view -F 0x400. Replicates were pooled by subsampling the same amount of reads per replicates using sambamba (v0.8.1)^58^ and samtools view (-s option). Peak calling was processed with MACS2 (https://github.com/macs3-project/MACS)^59^ (-q 0.01 --nomodel --extsize 147 --gsize=2.6e9) for H3K4me3, with MACS2 broad option for H3K27me3 (--broad -- broad-cutoff 0.1 -q 0.01 --nomodel --extsize 147 --gsize=2.6e9) and RECOGNICER^24^ (qvalue 0.001) /MACS2 (--broad --broad-cutoff 0.1 -q 0.01 --nomodel --extsize 147) for H2AK119ub1. SICER peak calling was performed using the following parameters -s XL -w 200 -rt 1 -f 150 -egf 0.74 -fdr 0.001 -g 600 -e 1000. Peaks overlap was defined using bedtools intersect having at least 1 bp of overlap between peaks. H2AK119ub1 signal was calibrated using reads mapping to the 601DNA and corresponding to the fully ubiquitylated H2AK119 nucleosome spiked-in in samples prior to ChIP (see ChIP M&M section). To evaluate difference in modifed histone ChIP-seq signal around the TSS of gene sets, Wilcoxon Mann-Whitney test was performed on the +/-1 kb window around the TSSs. ChIPSeeker^60^ and ChIPPeakAnno^61,62^ were used to annotate peaks on R^51^. The RegioneR^63^ package was used to perform peaks permutation and assess enrichment on USP21 sensitive gene features (*i.e.* TSS, enhancer, or gene body) compared to a random set of genes (using 10000 iterations of the resampleRegions function,, for Fig.3E, .3H, .S3C two right panels, .S3E, .S10G, .S10I) and to assess peak enrichment on genomic features (using 1000 iterations of the randomizeRegions function, for Fig 1G, 2K, 2M, S3C two left panels, S10H, S10K). Log10 (ChIP/Input) was computed using Deeptools with BamCompare using 50bp binsize. Heatmap and TSS plot were produced with computeMatrix and plotHeatmap or plotProfile. IGV was used for peaks and log10(ChIP/Input) signal visualisation. Heatmap were produced with computeMatrix using 50bp binsize +/-5kb of TSS or peak center. Heatmaps were generated with plotHeatmap function using unsupervised clustering (--hclust option), and metaplot using plotProfile function. IGV^64^ was used to visualize output signal tracks. Bedtools getfasta^65^ was used to extract FASTA sequences corresponding to peaks from a selected heatmap cluster. To identify transcription factors motif in the selected peak set, Homer^66^ was then used with the findMotifs.pl function and all other peaks of the heatmap as a background set. CpG islands and repeats elements were recovered from UCSC genome annotation database (https://hgdownload.soe.ucsc.edu/goldenPath/xenLae2/database/). For genome partitioning approach, chromHMM model was initialized using BinarizeBam on ChIP and Input sample file, using binsize of 150bp (-b 150). The model was trained using a random initiation (-init random), over binsize of 150bp (-b 150) and a maximum of 5000 iterations (-r 5000). SNPsplit was used for reads assignment to paternal and maternal copies as described in^19^.

## Data Availability

The *Xenopus laevis* ChIP-seq and RNA-seq data generated for this study will be referenced as PRJEB56442 at the European Genome-phenome Archive (EGA). Additional dataset used are from GSE73430 (Xenopus embryo RNA-seq), GSE125982 (sperm and replicated sperm H3K27me3 ChIP-seq), GSE126599 (Gastrulae H3K27me3,, and H3K4me3 ChIP-seq), and GSE75164 (sperm MBD-seq, spermatid H3K27me3), and GSE153531 (H2AK119ub1Cut&Run mouse embryos and gametes).

## Code Availability

All scripts used for these analyses are available on Github (https://github.com/FCValentin/H2AubPaper).

## AUTHORS CONTRIBUTION

Conceptualization: JJ; Methodology: JJ, VF-C, FB, LP; Validation: JJ, VF-C, FB; Formal analysis: VF-C, FB, MO, MG, VC; Investigation: JJ, VF-C, FB, NM, CF, JP, RG, MO, BH, KP, JGA; Writing original draft: JJ; Writing review and editing: JJ, FB, VFC. Visualisation: FB, VF-C; Supervision: JJ; Project administration: JJ; Funding acquisition: JJ.

## COMPETING INTERESTS

The authors declare no competing interests

## ACKOWLEDGEMENTS

J.J group is funded by ANR-21-CE13-0031-01 grant from the Agence Nationale de la Recherche and a Connect Talent grant from Ville de Nantes and Region pays de la Loire. M.O is funded by a postdoctoral fellowship from the Japan Society for the Promotion of Science (JSPS). R.G was supported for this work by a Career Development Award grant (CDA00019/2019-C) form the Human Frontier Science Program.. L.P group is supported by NIH award R24 OD031956. We would like to acknowledge the support of Xenbase.

## Notes

### Competing Interest Statement

The authors have declared no competing interest.

### Summary of Updates

Deposition of H2AK119ub1 by the polycomb repressive complexe-1 plays a key role in the initiation of facultative heterochromatin formation in somatic cells. Here we evaluate the contribution of sperm derived H2AK119ub1 to embryo development. In Xenopus laevis we found that H2AK119ub1 is present during spermiogenesis and into early embryonic development, highlighting its credential for a role in the transmission of epigenetic information from the sperm to the embryo. In vitro treatment of sperm with USP21, a H2AK119ub1 deubiquitylase, just prior to injection to egg, results in developmental defects associated with gene upregulation. Sperm H2AK119ub1 editing disrupts egg factor mediated paternal chromatin remodelling processes. It leads to post-replication accumulation of H2AK119ub1 on repeat element of the genome instead of CpG islands. This shift in post-replication H2AK119ub1 distribution triggered by sperm epigenome editing entails a loss of H2AK119ub1 from genes misregulated in embryos derived from USP21 treated sperm. We conclude that sperm derived H2AK119ub1 instructs egg factor mediated epigenetic remodelling of paternal chromatin and is required for embryonic development.

